## Supplemental Figures and Table for "SOX4 exerts contrasting regulatory effects on labor-associated gene promoters in myometrial cells"

**Supporting Information for**: SOX4 exerts contrasting regulatory effects on labor-associated gene promoters in myometrial cells.

Author List and Affiliations:

Nawrah Khader[1], Virlana M. Shchuka[1], Anna Dorogin[2,3], Oksana Shynlova[2, 3], and Jennifer A. Mitchell[1]

1. Department of Cell and Systems Biology, University of Toronto, Toronto, ON, M5S 3G5, Canada.
2. Lunenfeld Tanenbaum Research Institute, Sinai Health System, Toronto, ON, M5G 1X5, Canada.
3. Department of Obstetrics and Gynaecology, University of Toronto, ON, M5G 1E2, Canada.

Current address: Department of Cell and Systems Biology, University of Toronto, Toronto, Canada

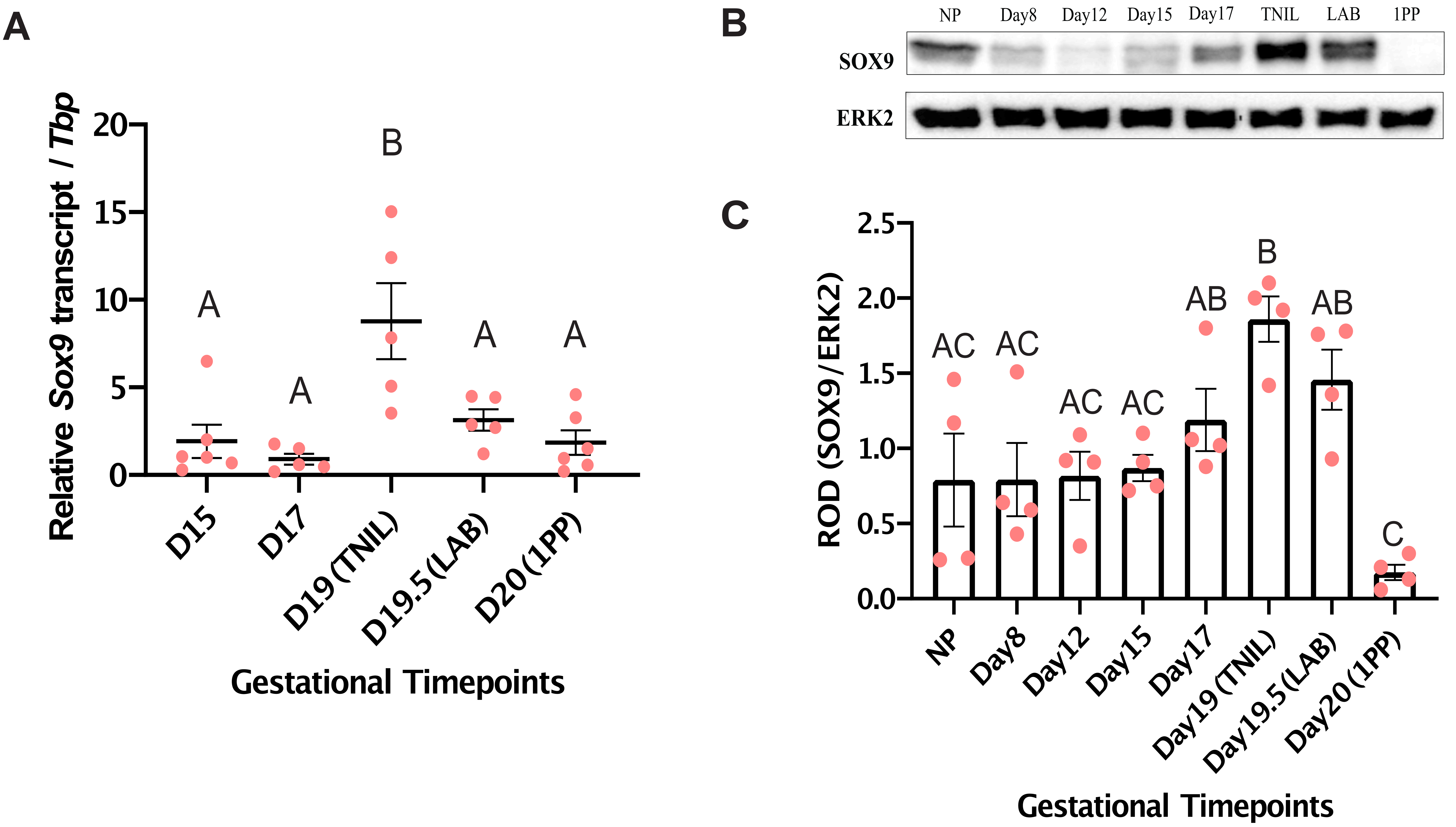


**SFig 1. SOX9 gene and protein expression is significantly elevated in murine term myometrium tissues.** (A) Transcript expression levels of *Sox9* was measured by RT-qPCR (±SEM) at indicated timepoints. Groups that exhibit significant differences ($p < 0.05$) as determined by one-way ANOVA are distinguished by different letters, while groups that do not show significant differences ($p > 0.05$) are labeled with the same letter.
(B) Representative immunoblot image of SOX9 and housekeeping protein ERK2 at gestational timepoints, indicated either by day (D) or status [term-not-in-labour (TNIL), labour (LAB), and postpartum (PP)].
(C) Densitometric analysis for SOX9 protein lysates extracted from mouse myometrium tissues at various gestational stages as indicated above. Groups that exhibit significant differences ($p < 0.05$) as determined by one-way ANOVA are distinguished by different letters, while groups that do not show significant differences ($p > 0.05$) are labeled with the same letter.

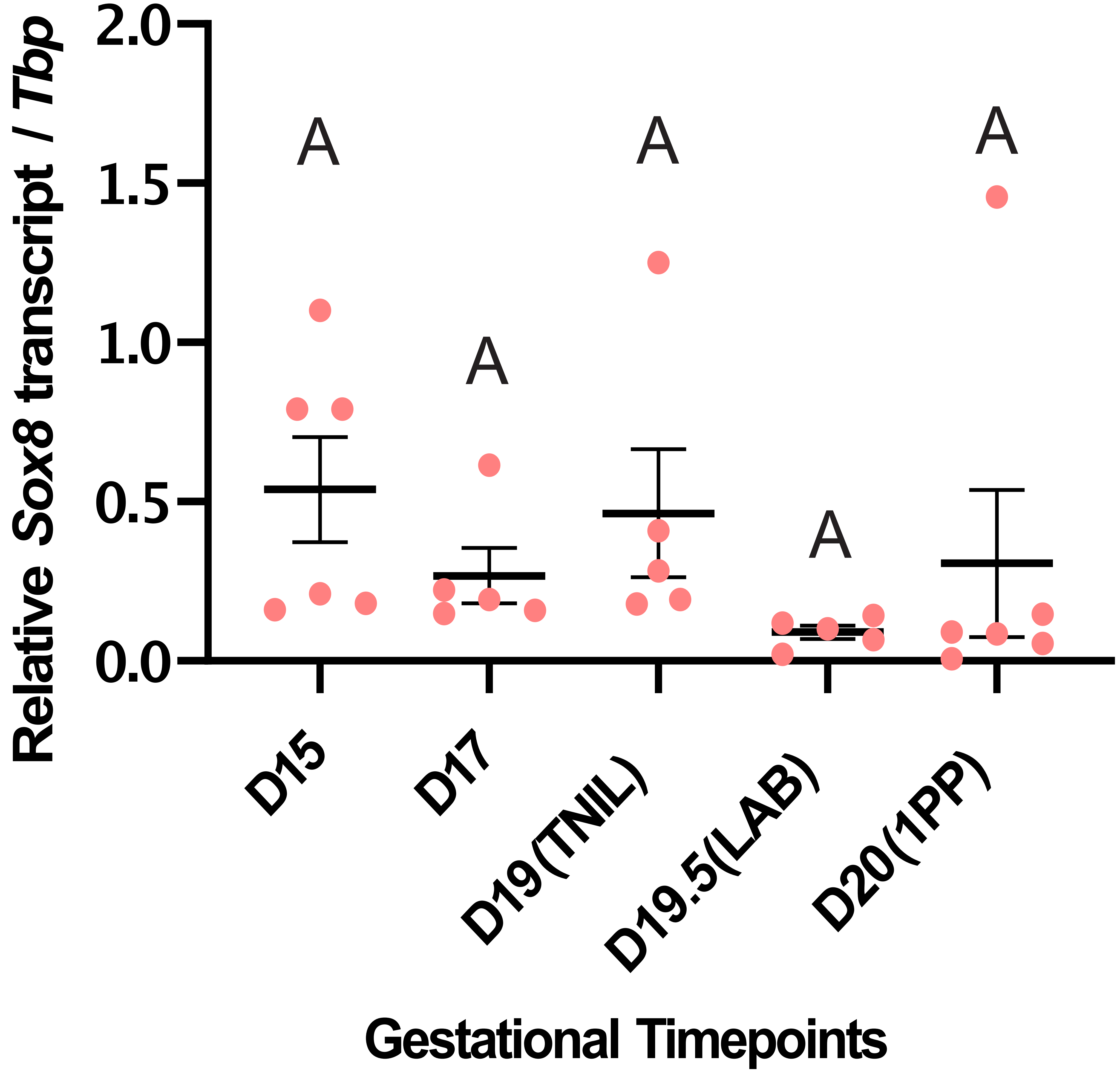


**SFig 2. The *Sox8* gene expression is not differentially expressed in the murine myometrium tissues across gestation.** Transcript levels of gene encoding the *Sox8* transcription factor, as measured by RT-qPCR (±SEM) at the designated timepoints.

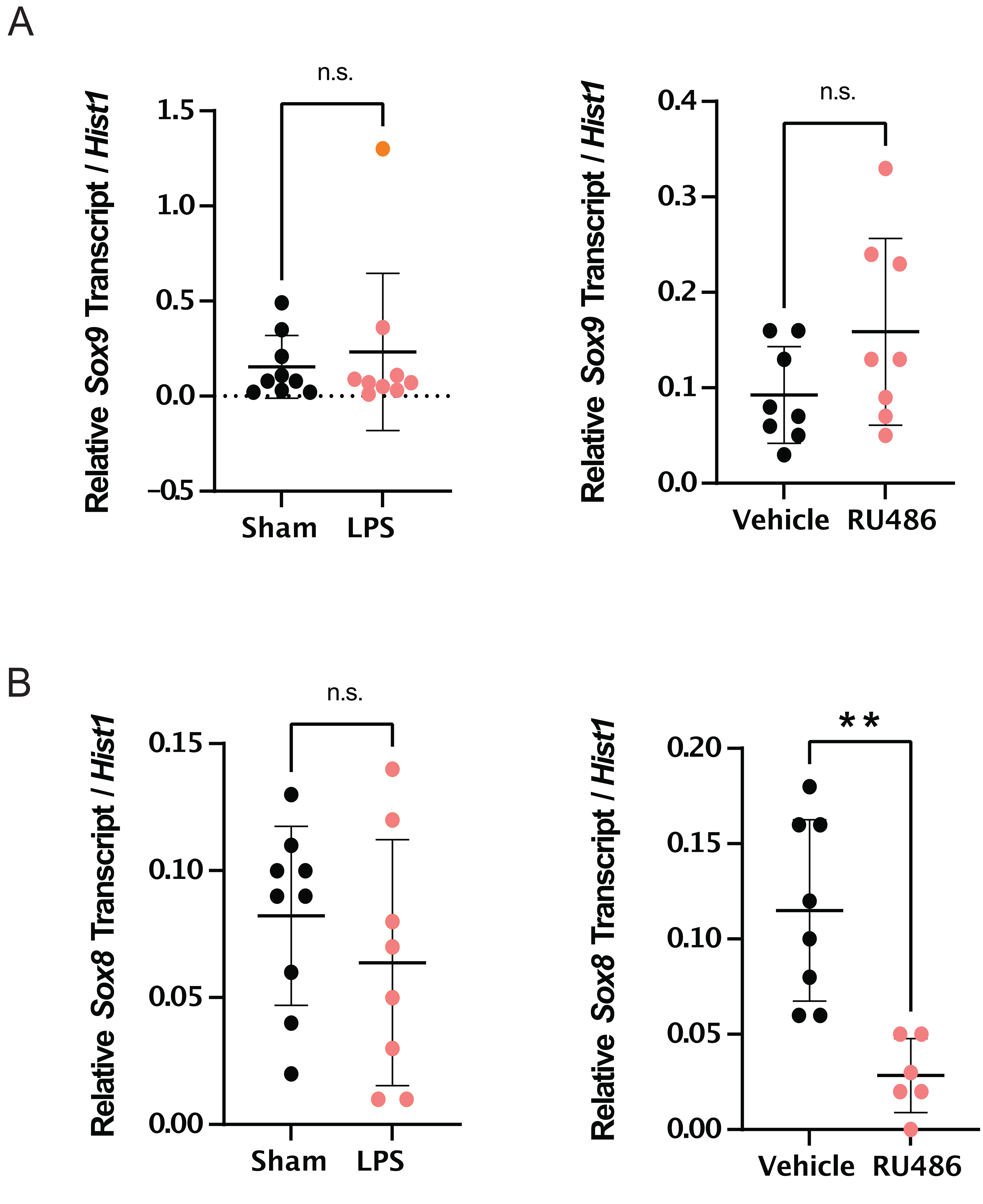


**SFig 3. *Sox8* is up-regulated during the onset of the RU486-mediated but not during LPS-induced preterm labor.** (A) Schematic of myometrium tissue collections at designated timepoints in two preterm labor induction models: local infection-simulation (LPS) and loss of progesterone (RU486).
Labour was induced by LPS (*left*) or RU486 (*right*), alongside sham and vehicle controls on day 15 and preterm labour occurred 18-24 hours post-injection as indicated by triangles. (B) Transcript levels of genes encoding the *Sox9* and *Sox8* transcription factors were measured by RT-qPCR (±SEM) at the timepoints corresponding to the schematic. Statistically significant differences in expression levels between control and preterm labor are marked by ** ($p < 0.01$).

**SUPPLEMENTAL TABLES**

**S1 Table. List of primers used in mouse labor-upregulated TF expression quantification experiments.**

| TF Transcript Target | Forward Primer 5’->3’ | Reverse Primer 5’->3’ | Size (bp) |
|---|---|---|---|
| *Sox4* | TTGCCGACTTCACCTTCTTTC | GACAAGATTCCGTTCATCCAGC | 104 |
| *Sox7* | CCGACCTTCAGGGGACAAGA | ATCTTGCTGAGCTCCGCGTT | 135 |
| *Sox8* | CCGGCCAGTCTTCACACTCT | GCGAGAAGAGGCCCGTTTGTG | 113 |
| *Sox9* | AGCAGCGACGTCATCTCCAA | GCTGCTTCGACATCCACACG | 184 |
| *Tbp* | CTCAGTTACAGGTGGCAGCA | ACCAACAATCACCAACAGCA | 186 |
| *Hist1* | GGCCAAGGCTTCCAAGAAGT | CCACCTTGTAGTGGCTCTTGATA | 137 |

**S2 Table. List of primers used to clone labor-associated gene promoters into target reporter vectors for luciferase assays.**

| Promoter | Forward Primer 5'->3' | Reverse Primer 5'->3' | Size (bp) |
|---|---|---|---|
| *Gja1* | TCTCCTGAAGGAATGACCCATCCA | gtctgggcacctcTCTTTCACTTAATGAA AGTGAAGCC | 503 |
| *Gja1* (cloning) | gaggatatcaagatctTCTCCTGAAGGA ATGACCCATCCA | ttggcatcttccatggGTCTGGGCACCTC TCTTTCACT | n/a |
| *Fos* | gaggatatcaagatctACTTATTTACAA TCCTTCACTTGCT | ttggcatcttccatggGGTCGAAGTTTGG GGAAAGCC | 901 |
| *Ptgs2* | AGCATTCCGATGAAGTGGAGCT | GGAGGTGGCAGTAGTGGTGG | 757 |
| *Ptgs2 (cloning)* | gaggatatcaagatctAGCATTCCGATG AAGTGGAGCT | ttggcatcttccatggGGAGGTGGCAGT AGTGGTGG | 789 |
| *Mmp11* | TGCCAAGTGTCAGTAGAGGTCAG | CTGCTGGGCCTGCTGGG | 608 |
| *Mmp11 (cloning)* | gaggatatcaagatctTGTGCCAAGTGT CAGTAGAGGT | ttggcatcttccatggTGCTGGGCCTGCT GGG | n/a |
